# Coding of chromatic spatial contrast by macaque V1 neurons

**DOI:** 10.1101/2021.02.12.430975

**Authors:** Abhishek De, Gregory D. Horwitz

## Abstract

Color perception relies on spatial comparisons of chromatic signals, but how the brain performs these comparisons is poorly understood. Here, we show that many V1 neurons compare signals across their receptive fields (RF) using a common principle. We estimated the spatial-chromatic RFs of each neuron, and then measured neural selectivity to customized colored edges that were sampled using a novel closed-loop technique. We found that many double-opponent (DO) cells, which have spatially and chromatically opponent RFs, responded to chromatic contrast as a weighted sum, akin to how simple cells respond to luminance contrast. Other neurons integrated chromatic signals non-linearly, confirming that linear signal integration is not an obligate property of V1 neurons. The functional similarity of DO and simple cells suggests a common underlying neural circuitry, promotes the construction of image-computable models for full-color image representation, and sheds new light on V1 complex cells.

## INTRODUCTION

Color perception depends on the spectral composition and spatial organization of lights in a visual scene (Monnier and Shevell, 2003; Shevell and Monnier, 2005). Spectral signals across space are thought to contribute to scene segmentation as well as facilitating object recognition and enhancing visual memory (Wurm et al., 1993; Fine et al., 2003; Spence et al., 2006). Despite its importance for visual function, the spatial integration of spectral signals by neurons in the visual system is poorly understood.

The first stage of the primate visual system at which spatial comparison of spectral signals is implemented by individual neurons is area V1; many neurons in V1 are tuned for orientation and spectral properties of edges. Simple and double-opponent (DO) cells are tuned for the orientation of luminance and chromatic edges, respectively (Ringach, 2002; Conway and Livingstone, 2006; Johnson et al., 2008; De and Horwitz, 2020). Simple cells combine signals from the L- and M-cone photoreceptors with the same sign and are therefore cone non-opponent. They respond to light increments in one part of their receptive fields (RFs) and decrements in another, rendering them sensitive to spatial luminance contrast. DO cells are spatially opponent, like simple cells are, but they combine signals from at least two different cone photoreceptor types antagonistically within individual RF subfields (Daw, 1968; Conway, 2001; De and Horwitz, 2020).

Simple cells combine signals across their RFs approximately as a weighted sum (Hubel and Wiesel, 1959; Movshon et al., 1978b; Carandini et al., 1997). These neurons can therefore be thought of as linear filters that operate on the retinal image. This insight has propelled scientific progress in at least three ways. First, it facilitated the construction of image-computable models of achromatic image representation (Marr and Hildreth, 1980; Adelson and Bergen, 1991). Second, it shed light on cortical circuitry: simple cells receive excitation (“push”) and inhibition (“pull”) with opposite spatial tuning, which appears to be a critical step for establishing linearity in the face of nonlinear input from the lateral geniculate nucleus (Ferster, 1988; Hirsch et al., 1998; Ferster and Miller, 2000). Third, it provided an essential building block for quantitative models of V1 complex cells (Movshon et al., 1978a; Adelson and Bergen, 1985), neurons in higher-order cortical areas (Simoncelli and Heeger, 1998; Cadieu et al., 2007; Freeman et al., 2013), and a variety of psychophysical phenomena (Adelson and Bergen, 1985; Beaudot and Mullen, 2006; Graham, 2011). Currently, all of these advances have been restricted to the achromatic domain. A similar quantitative understanding of image representation in the chromatic domain is severely lagging and is necessary to extend these advances to natural, full-color images.

The nature of the spatial antagonism implemented by DO cells has important implications for their contributions to vision. For example, consider a hypothetical DO cell that compares cone-opponent signals between the left and right halves of its RF (**Figure 1A**). If the cell integrates signals linearly and with equal weight, then it is well suited for extracting vertical chromatic edges (**Figure 1B–D**). In this case, the excitation due to a preferred light in one half of the RF can be cancelled by the same light appearing in the neighboring half. On the other hand, if the cell integrates signals nonlinearly, for example, by weighting contrast increments more heavily than contrast decrements, it would encode both edges and surfaces (**Figure 1C,F**). In this case, a light in one half of the RF would fail to cancel the same light appearing in the neighboring half.

**Figure 1.**
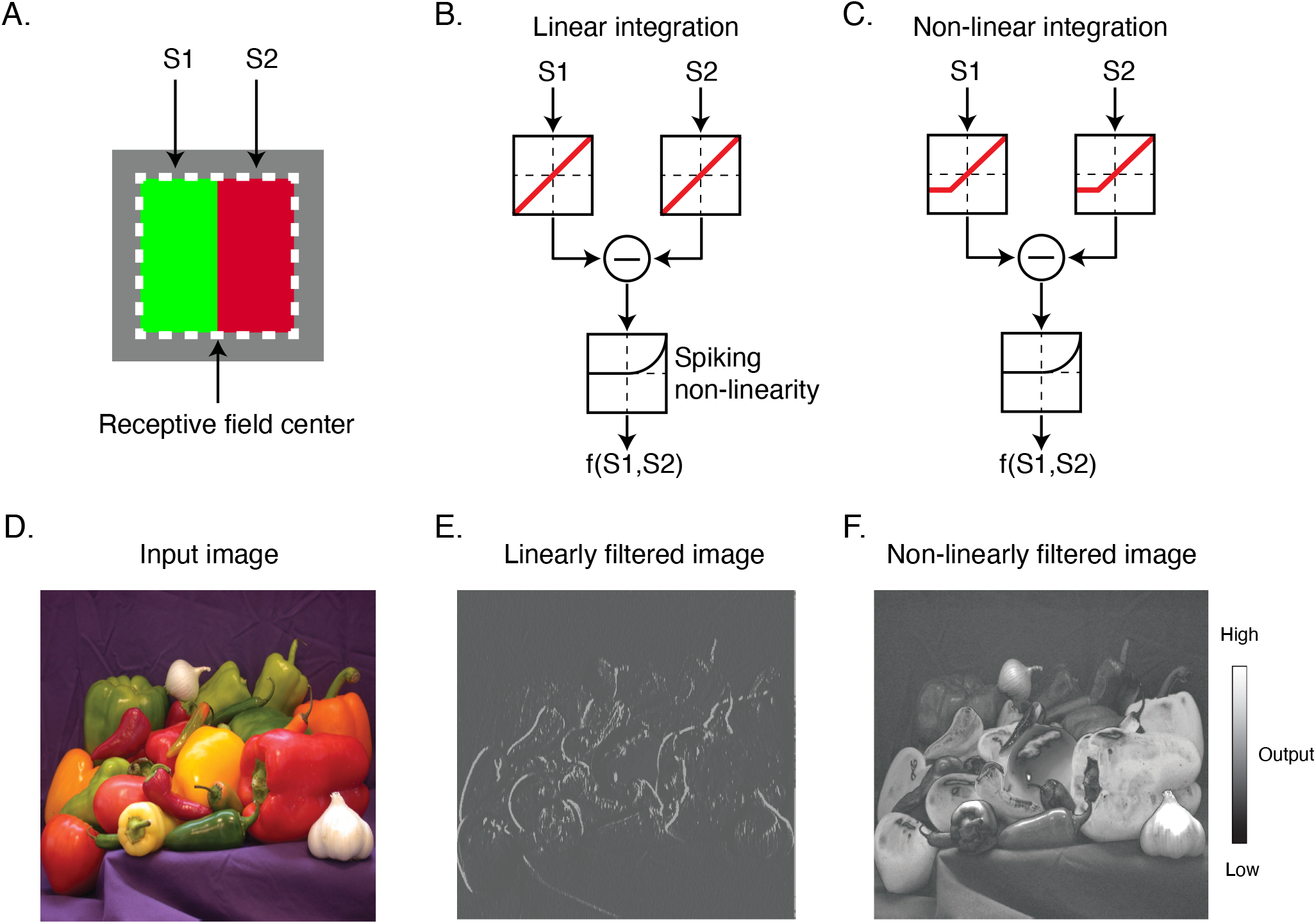
Linear and non-linear image filtering **A.** A hypothetical V1 DO cell that is excited by a red light on one side of its receptive field and a green light on the other. S1 and S2 represent stimulation of the two subfields of the receptive field. **B.** A linear spatial filter that sums the stimulation of each subfield and generates a response via a spiking non-linearity. Note that the drive from each of the subfields is combined linearly before being transformed by the spiking non-linearity. **C.** A non-linear spatial filter that partially rectifies S1 and S2 prior to summation. **D.** An example natural image. **E.** Output of the linear spatial filter of the input image. **F.** Same as **E** but for the non-linear filter.

In this study, we measured the spatial integration of visual signals by individual V1 neurons in awake, fixating monkeys. Neurons can combine signals in many ways, but we focus on linear combinations because of their theoretical significance. Linearity is a mathematical ideal that is never fully realized by neurons, so we do not ask whether neurons integrate visual information across their RFs linearly but instead quantify how well a linear model describes spatial integration relative to a more flexible model. Models were fit to data collected using a closed-loop stimulus generation technique (Bölinger and Gollisch, 2012; Horwitz and Hass, 2012). The closed-loop technique involved the online construction of optimally oriented and positioned edges that varied in color but drove identical spike count responses. An advantage of this approach is that it is robust to output nonlinearities that complicate other approaches (DeAngelis et al., 1993; Gollisch and Herz, 2012). It also avoids the necessity of adopting a single definition of stimulus contrast, which is particularly important when probing responses to chromatic stimuli because no universally accepted definition of contrast applies to all color directions and how linear a neuron appears depends on stimulus contrast (Albrecht and Hamilton, 1982; Mullen, 1985).

Using this technique, we found that many DO cells responded linearly to differences in cone-opponent signals across their RFs, in quantitive similarity to how simple cells respond to spatial differences in luminance. Other V1 neurons combined signals non-linearly, confirming the existence of non-linear, cone-opponent neurons in V1 and the ability of the closed-loop technique to detect them. These results suggest that DO cells, like simple cells, are building blocks for complex cells, a proposal that explains several previous results and provides new insight into the functional role of color-sensitive V1 complex cells.

## RESULTS

We analyzed the spiking responses of 98 V1 neurons in two awake, fixating male macaque monkeys (69 from Monkey 1, 29 from Monkey 2). Each neuron was stimulated with white noise, and data were analyzed by spike-triggered averaging to identify a pair of functionally distinct subfields within the classical RF. Visual stimulation was then targeted to these subfields to characterize their individual and joint impact on neuron’s firing rate.

### RF characterization

Spike-triggered averages (STAs) of some neurons resembled uniform blobs, indicating consistent spectral sensitivity across the RF. Other STAs were spatially structured, for example, consisting of a set of yellow pixels adjacent to a set of blue pixels (**Figure 2A**). These structured STAs are a signature of neurons that compare spectral information across space. These neurons only were tested; if an STA did not reveal at least two distinct subfields, the neuron was passed over for data collection.

**Figure 2.**
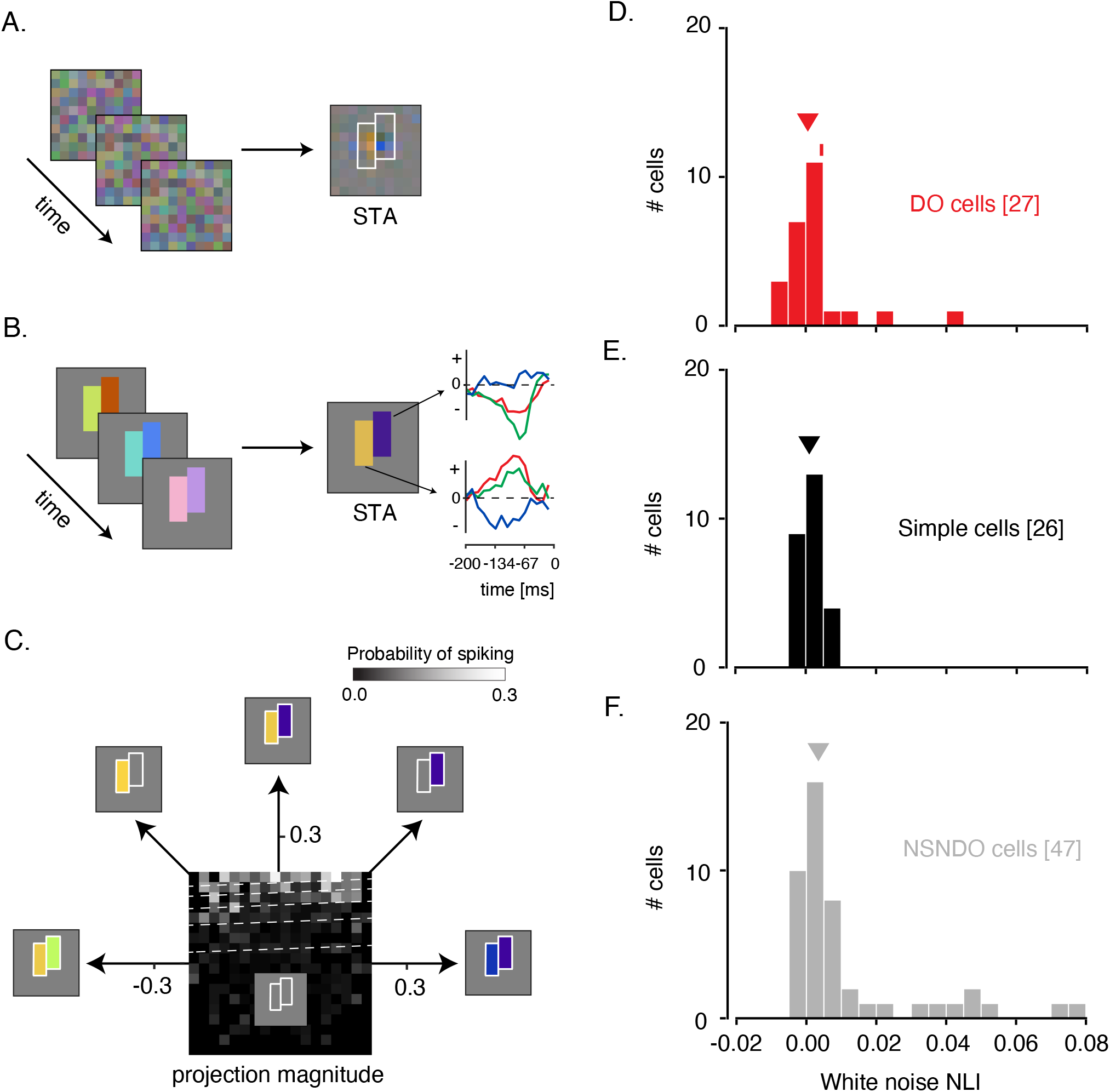
White noise analysis of RF structure and spatial integration **A.** Checkerboard white noise stimulus (left), spike-triggered average (STA; right). Two sets of contiguous pixels were yoked to create two hyperpixels, each of which stimulated one RF subfield (white outlines). **B.** The customized white noise stimulus (left) and STA at the peak frame (middle). Red, green, and blue curves (right) represent the average red, green, and blue phosphor intensities, relative to background, as a function of time before a spike. **C.** A firing rate map for the example DO cell. The probability of spiking (gray scale) is plotted as a function of projection magnitudes of the stimulus onto the right and left halves of the STA (along the 45 and 135 directions, respectively). Dashed white lines are contours of constant spiking probability from a GLM fit to the data. **D.** Histogram of white noise NLIs for DO cells. The NLI of the example neuron is marked with a tick, and the median is marked with a triangle **E.** Same as **D** but for simple cells. **F.** Same as **D** but for the cells that were neither simple nor DO.

A major goal of these experiments was to characterize spatial integration within the RFs of V1 neurons. The white noise checkerboard stimulus is ill-suited for achieving this particular goal because it modulates V1 neurons weakly, and nonlinear spatial integration might appear linear in response to small perturbations. To drive the neurons more effectively, we customized the stimulus to each neuron studied. From the STA, we identified two contiguous groups of pixels, each covering a single RF subfield, and yoked each group into a “hyperpixel”. White noise modulation of the two hyperpixels stimulated the two subfields strongly and independently, driving a wide range of firing rates (*Phase 2* of the protocol, **see Experimental Protocol**, **Figure 2B & Figure S1**).

Neurons with spatially opponent RFs were classified into three categories on the basis of their responses to the hyperpixel white noise stimulus. Twenty-six neurons were classified as simple cells, 27 were classified as DO cells, and 47 neurons were neither simple nor DO and were classified conservatively as “NSNDO” (neither simple nor double-opponent). For each neuron studied, the colors on the two sides of the hyperpixel STA were complementary or nearly so (Pearson’s r between the two sides: −0.94 ± 0.11 (mean ± SD) for simple cells, −0.76 ± 0.23 for DO cells, and −0.77 ± 0.24 for NSNDO cells). Many neurons in the NSNDO category are likely to be complex cells although a decisive classification would require an analysis of F1/F0 response amplitude modulation which we did not measure (Skottun et al., 1991). Some NSNDO neurons might have been classified as DO in other studies, an issue to which we return in the Discussion.

### Measuring spatial integration using white noise

We quantified interactions between RF subfields using an approach similar to one used previously to study interactions between the stimulus features that trigger spikes in complex cells (Touryan et al., 2002; Rust et al., 2005). In these previous studies, white noise stimuli were projected onto the plane spanned by the first and second principal components of the stimuli the drove spikes. Similarly, we projected the hyperpixel white noise stimuli onto the two halves of the STA (**see White noise analysis of signal combination across subunits**). These two projection values reveal how similar the stimulus was to the two halves of the STA; the larger the projection value, the more of the STA is present in the stimulus. We visualized a firing rate map by binning stimulus projections and calculating the proportion of stimuli in each bin that drove a spike (**Figure 2C)**. The probability of spiking increased with the stimulus projection onto individual RF subfields, and it rose more steeply when both projections increased together.

To analyze spatial integration between RF subfields, we fit the data with a generalized linear model (GLM) and a generalized quadratic model (GQM) (**see White noise analysis of signal combination across subunits; Figure S2A**). We then quantified the ability of the two models to classify stimuli as spike-triggering or not using ROC analysis (**Figure S2B**) (Green and Swets, 1966). Classification error rates of the GLM and GQM were summarized with a white noise non-linearity index (NLI) (**see White noise non-linearity index**). A white noise NLI < 0 indicates that the GLM provides more accurate predictions than the GQM, and an NLI > 0 indicates that the GQM provides more accurate predictions than the GLM. An NLI of 0 occurs if the GLM and GQM make identical predictions, which can occur because the GLM is a special case of the GQM with three parameters set to zero. Because of these extra parameters, the GQM always fits the training data as well or better than the GLM. To compare the two models fairly, we tested the model on data that had been held out from the fitting using 10-fold crossvalidation (Browne, 2000).

NLIs differed across the three cell types (median white noise NLI for simple cells = 0.0009, DO cells = 0.0005, NSNDO cells = 0.0034; p=0.02, Kruskal-Wallis test; **Figure 2D–F**). Comparison between simple and DO cells revealed no significant difference between them (p=0.99, Mann-Whitney U test). To the contrary, NLIs of simple and DO cells were both lower than those of the NSNDO cells (p<0.05, Mann-Whitney U tests). We conclude that simple and DO cells are similarly linear and are more so than other V1 neurons that also have spatially structured STAs.

In interpreting these data, it is important to note that a lack of evidence for a difference between simple and DO cells is not evidence that a difference does not exist. This experiment probed neurons with low-contrast, rapidly modulated stimuli (**Figure S1**). The possibility remains that differences between simple and DO cells become evident when they are tested with stimuli of higher contrast or longer duration. We tested this possibility in *Phase 3* of our experimental protocol, as described below.

### Measuring spatial integration using the isoresponse method

For each neuron, we found a collection of stimuli that evoked the same response. Each stimulus was spatially identical to the hyperpixel STA, but the contrast of the two hyperpixels was adjusted according to the algorithm described in **Contrast staircase procedure**. Negative contrasts were allowed.

To appreciate the necessity of this technique, it is useful to consider a classical alternative. A classic test of linearity is to present one stimulus at the receptive field of a recorded neuron, then another, and then both together. If the response to the combined stimulus does not equal the sum of responses to the two components, the neuron is not linear. However, this test is sensitive to nonlinearities that are logically distinct from the linearity of spatial summation and are present in otherwise linear cortical neurons (e.g. spike firing thresholds and saturating contrast-response functions). An alternative approach is to find a collection of stimuli that evoke the same response from an isolated neuron and analyze these stimuli to identify the features they share. This approach has been used previously to study signal integration in the salamander retina and locust auditory receptor cells (Gollisch et al., 2002; Bölinger and Gollisch, 2012). It has also been used previously in macaque V1 to analyze the linearity of signal integration across cone types by individual neurons (Horwitz and Hass, 2012), but it has not been used previously to analyze the linearity of signal integration across a V1 RF.

If a neuron combines cone-contrast signals linearly across its RF, then stimuli that drive the same response will lie on lines when represented in the stimulus space shown in **Figure 2C**. If the stimuli lie on a curve instead of a line, the hypothesis of linear spatial summation can be rejected. This approach makes no assumptions about static output nonlinearities downstream of spatial integration whereas the GLM and GQM assumed a logistic function. It also does not assume linearity of cone signal integration within individual RF subfields. This assumption was needed to reduce the RGB values that identify each stimulus frame to two stimulus projections for the GLM and GQM analyses.

In **Figure 2C**, each point represents a stimulus, distance from the origin represents contrast relative to the background, and angle represents contrast between the two sides of the stimulus. Within this plane, a search was performed to find physically distinct stimuli that evoked the same neuronal response. Angles were selected pseudo-randomly, and distances were titrated by a staircase procedure until a target firing rate was achieved (**Figure S3**). To mitigate the impact of spontaneous spiking activity on staircase procedure, target firing rates were well above baseline firing rates (95/98 neurons had target firing rates that were greater than the 95^th^ percentile value of their respective baseline firing rate distribution). Target firing rates did not differ across cell types (p=0.57, Kruskal-Wallis Test).

For some neurons, staircase termination points lay close to a line when plotted in the stimulus space (**Figure 3A–B**). This result shows that the excitation produced by a preferred light at one part of the RF can be cancelled by an anti-preferred light at a neighboring part with a fixed constant of proportionality over the entire gamut of our video display. This cancellation is consistent with linearity of spatial integration (**Figure 1A**) and not with differential sensitivity to contrast increments and decrements (**Figure 1B**). However, not all neurons behaved this way. For some neurons, staircase termination points lay on a curve (**Figure 3C**), indicating nonlinear spatial integration.

**Figure 3.**
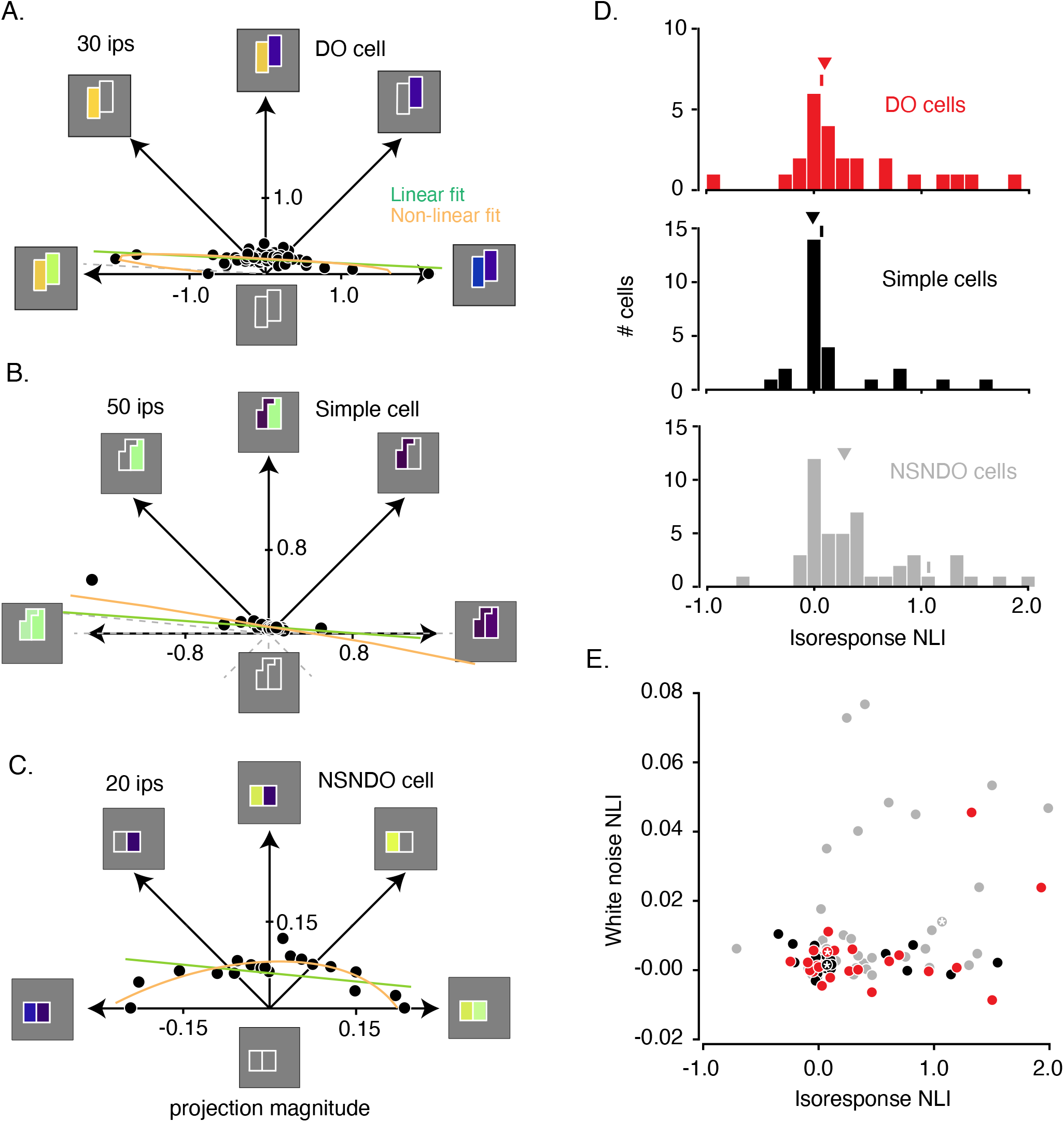
Analysis of isoresponse contours. **A.** Data from the example DO cell shown in **Figures 2A–C**. Dots indicate staircase terminations (target firing rate = 30 ips) and gray dashed lines indicate staircases that exceeded the monitor gamut. Linear (green) and non-linear (orange) fits to the data are similar. **B.** Same as **A** but for a simple cell (target firing rate = 50 ips). L+M spectral sensitivity manifests as bright green (ON) or dark purple (OFF) when probed with the RGB white noise (Chichilnisky and Kalmar, 2002). **C.** Same as **A** but for a cell that was neither simple nor DO (target firing rate = 20 ips). **D.** Histogram of isoresponse NLIs. NLIs of example neurons are marked with ticks, and medians are marked with triangles. **E.** Scatter plot of isoresponse NLIs and white noise NLIs. Example neurons are marked with white asterisks.

To determine quantitatively whether a line or a curve provided the better description of the staircase termination points for each neuron, we compared linear and quadratic models fits (**see Evaluating model fits to staircase termination points; Figure S4**). We defined an isoresponse non-linearity index (isoresponse NLI) similarly to the white noise NLI defined previously (**see Isoresponse non-linearity index**). An isoresponse NLI of 0 indicates that the linear and quadratic models made equally accurate response predictions, NLI < 0 indicates that the linear model predicts responses more accurately than the quadratic model, and NLI > 0 indicates that the quadratic model predicts responses more accurately than the linear model. Cross validation ensured that the quadratic model does not achieve greater prediction accuracy simply by virtue of having more parameters.

NLIs of simple cells and DO cells were close to zero and did not differ significantly (median isoresponse NLI for DO cells = 0.1007, median isoresponse NLI for simple cells = −0.0097; p=0.14, Mann-Whitney U test; **Figure 3D**). In contrast, NLIs were greater for the NSNDO neurons (median isoresponse NLI = 0.2822, p=0.02, Kruskal-Wallis test). We conclude that simple and DO cells are similarly linear over the range that we were able to test given the limits of our display, and that they are more linear than other neurons in V1.

Despite the many methodological differences between *Phases 2* and *3* of the experiment, the results were similar for individual neurons (**Figure 3E**). Isoresponse NLIs were positively correlated with the white noise NLIs (r = 0.30, p = 0.001, Spearman’s correlation between isoresponse NLI and white noise NLI). This correlation was driven primarily by NSNDO cells (r = 0.41, p = 0.004, Spearman’s rank correlation) and not by DO (r = 0.01, p = 0.95, Spearman’s rank correlation) or simple cells (r = −0.19, p = 0.34, Spearman’s rank correlation). We conclude that some nonlinearities that manifest in responses to white noise also manifest in responses to more classical stimuli, and that nonlinearities that were not manifested in responses to white noise were similarly common among simple and DO cells.

## DISCUSSION

A fundamental goal of visual neuroscience is to characterize the transformation from light stimulation of individual neurons to spiking responses. In this study, we characterized the spatial integration by individual V1 neurons using a combination of white noise RF mapping, closed-loop isoresponse measurements, and statistical model comparisons. We found that DO cells integrate signals across their RFs roughly linearly, like simple cells and in contrast to other V1 neurons, which were less linear on average. Below, we compare our results to those of previous studies and discuss the impact of our cell classification criteria on the results. We then discuss the implications of our results on the circuitry that underlies DO and simple cells and how each cell type may contribute to downstream image processing. We conclude with speculations on parallels between the processing of color and other stimulus features in V1 by complex cells.

### Relationship to previous work

Linearity in the visual system is “a rare and (apparently) prized commodity in neural signal processing” (Shapley, 2009). The linearity of V1 simple cells is not an accident of random convergence of LGN afferents but rather the product of specialized excitatory and inhibitory circuitry (Ferster, 1988; Tolhurst and Dean, 1990; Hirsch et al., 1998). The discovery that V1 simple cells combine signals linearly across their RFs contributed to scientific progress in many ways. It provided a valuable bridge between neurophysiology and the fields of psychophysics and computer vision. It provided guidance for how to characterize neuronal stimulus tuning efficiently. It served a basis for more elaborate models; all V1 neurons exhibit some degree of nonlinearity, but the linear model remains a cornerstone of even nonlinear V1 models (Carandini et al., 2005; Carandini, 2006).

Color can be quantified in many ways. In this study, we used the intensity of individual (display-specific) phosphor channels relative to the background. The fact that DO cells and simple cells combined light intensities across space approximately linearly in this representation shows that it is a reasonable one for analyzing V1 neurons. We note that any color representation that is a linear transformation of this device-specific color space would also have this property (e.g. cone-contrast, cone-excitation-differences-from-background, Derrington-Krauskopf-Lennie, etc).

Nearly all quantitative studies of V1 RFs have used achromatic stimuli. Extending quantitative RF mapping to color is complicated by the curse of dimensionality. As the dimensionality of the color space increases from 1-D (achromatic) to 3-D (full color), the number of possible spatial combinations grows exponentially. Classic workarounds include the use of gratings, which have a highly constrained spatial structure and/or cone isolating stimuli, which are most useful for analyzing neurons that combine signals linearly across cone type. Our solution was to map the RF of each neuron with white noise and then customize spatiochromatic patterns on the basis of these maps.

Two previous studies investigated spatial integration by DO cells. Using the 2-bar interaction technique, Conway et al. (2002) found that most color-sensitive V1 neurons responded maximally when a pair of different cone-isolating bars appeared side-by-side inside the RF. This maximal response exceeded the response to either bar in isolation, consistent with linearity as well as with other models. Conway and Livingstone (2006) measured the responses of DO neurons to cone-isolating stimuli at individual RF locations. They assumed that the excitatory response to a contrast increment had the same magnitude, but opposite sign, as the suppression to a contrast decrement. This need not be the case; retinal and LGN ON and OFF pathways are asymmetric, and some cone-opponent pathways particularly so (De Valois et al., 2000; Chichilnisky and Kalmar, 2002; Chatterjee and Callaway, 2003; Tailby et al., 2008). Nevertheless, most of the DO cells they studied showed clear signs of push-pull inhibition, which is consistent with linearity (Tolhurst and Dean, 1990; Ferster, 1994; Hirsch et al., 1998; Ferster and Miller, 2000). Our findings extend these results by demonstrating the linearity of spatial integration directly through simultaneous stimulation of functionally distinct RF subfields, and they suggest that the assumption made by Conway and Livingstone (2006) is a reasonable one for V1 DO cells.

### Cell categorization criteria

We classified simple and DO cells on the basis of their responses to the hyperpixel white noise stimulus. Specifically, we segregated simple from DO cells on the basis of cone weights derived from the STA. By definition, simple cells have large, non-opponent L- and M-cone weights, and DO cells have cone-opponent weights. We used the same cone weight criteria that we used previously to facilitate comparison between studies (De and Horwitz, 2020).

The cone weight criteria for inclusion into the simple cell and DO cell categories were asymmetric for two reasons. First, cone non-opponent V1 cells (e.g. simple cells) typically have smaller S-cone weights than do cone-opponent cells (e.g. DO cells) (Johnson et al., 2004; Horwitz et al., 2007). Second, the variability in estimated L- and M-cone weights is greater for non-opponent cells than opponent cells (Horwitz et al., 2007). Reclassifying cells with different criteria did not change the main results of this study (**Figure S5**).

Most other recent studies of DO cells used cone-isolating stimuli, which cannot reveal interactions among cone types (Conway, 2001; Johnson et al., 2001, 2004; Conway and Livingstone, 2006; Johnson et al., 2008). In contrast, we used a stimulus set that modulated all three cone types together in a variety of proportions. Nonlinear interactions between cone types complicate the interpretation of RF maps that are separated by cone type. In further distinction from other studies, we stimulated DO cells with colored edges to confirm the spatial and spectral sensitivity inferred from the STAs.

We classified neurons with nonlinear responses to white noise as NSNDO. This criterion was necessary to satisfy the assumptions underlying the conversion of the STA to cone weights. Importantly, this criterion did not force the result of linearity in DO cells. Spike-triggered covariance (STC)—the technique we used to detect nonlinearities (**see Spike-triggered covariance analysis**)—detects only a subset of nonlinearities, and nonlinearities that are clear with high-contrast, long-duration stimuli are not always detectable with white noise (Tanabe and Cumming, 2008). Nevertheless, we found that nonlinearities detected in *Phase* 2 of our experiment were a good indicator of nonlinearity over the greater stimulus duration and range of contrasts in *Phase 3*, principally for the NSNDO cells (**Figure 3E**).

We speculate that some cells that we classified as NSNDO on the basis of nonlinearities in responses to white noise would have been classified as DO in other studies (Conway and Livingstone, 2006; Johnson et al., 2008). Whether these neurons are more usefully classified as nonlinear DO cells, partially rectified complex cells, or something else entirely is an important question that is partly physiological and partly semantic. In any case, the major finding of this study is that a population of DO cells combines cone-opponent signals across their RFs approximately as linearly as simple cells combine non-opponent signals, a result that stands despite the existence of other V1 cells with nonlinear spatial integration.

### DO and simple cells: Neural circuitry

Spatial linearity of V1 simple cells is based on excitatory and inhibitory pools of LGN afferents that carry distinct signals (Ferster, 1988; Tolhurst and Dean, 1990; Hirsch et al., 1998). The spectral sensitivity of a V1 neuron is determined by the afferents that contribute to each of these pools. Pooling LGN afferents with the same sign (ON or OFF) creates non-opponent spectral sensitivity. Simple cells are excited by L-ON and M-ON afferents (and inhibited by L-OFF and M-OFF) in one part of their RFs and have the reverse tuning in another part. In contrast, pooling the same afferent signals to produce cone-opponency (e.g. L-OFF with M-ON) with otherwise identical circuitry would produce a DO cell. This may be the primary difference between simple and DO cells. The implementation of the spatial differencing operation may be common to both cell types.

Simple cells are thought to provide the dominant input to complex cells (Hubel and Wiesel, 1962). Under a standard model, complex cells pool signals from simple cells with overlapping RFs and common preferred orientation (**Figure 4A**). One possibility is that some complex cells also receive input from DO cells (**Figure 4B**). We speculate that complex cells receiving simple cell input only are luminance-sensitive **(Figure 4C)** whereas those that receive input from simple and DO cells are both color-and luminance-sensitive **(Figure 4D)**. This conjecture is consistent with the observations that the preferred orientation of color-sensitive complex cells is maintained across color directions (Johnson et al., 2001). It is also consistent with the observation that color-sensitive complex cells have multiple preferred color directions by STC analysis (Horwitz et al., 2007) and have no null directions in cone-contrast space (Horwitz and Hass, 2012). An alternative possibility is that the spectral sensitivity of color-sensitive complex cells arises entirely from their DO cell inputs (Michael, 1978), which have been reported to be similarly responsive to chromatic and luminance contrast (Johnson et al., 2001, 2004, 2008). In either case, DO cells are a likely basis for the chromatic sensitivity of color-sensitive complex cells (Michael, 1978). Measuring functional connectivity between DO and complex cells could test this hypothesis.

**Figure 4.**
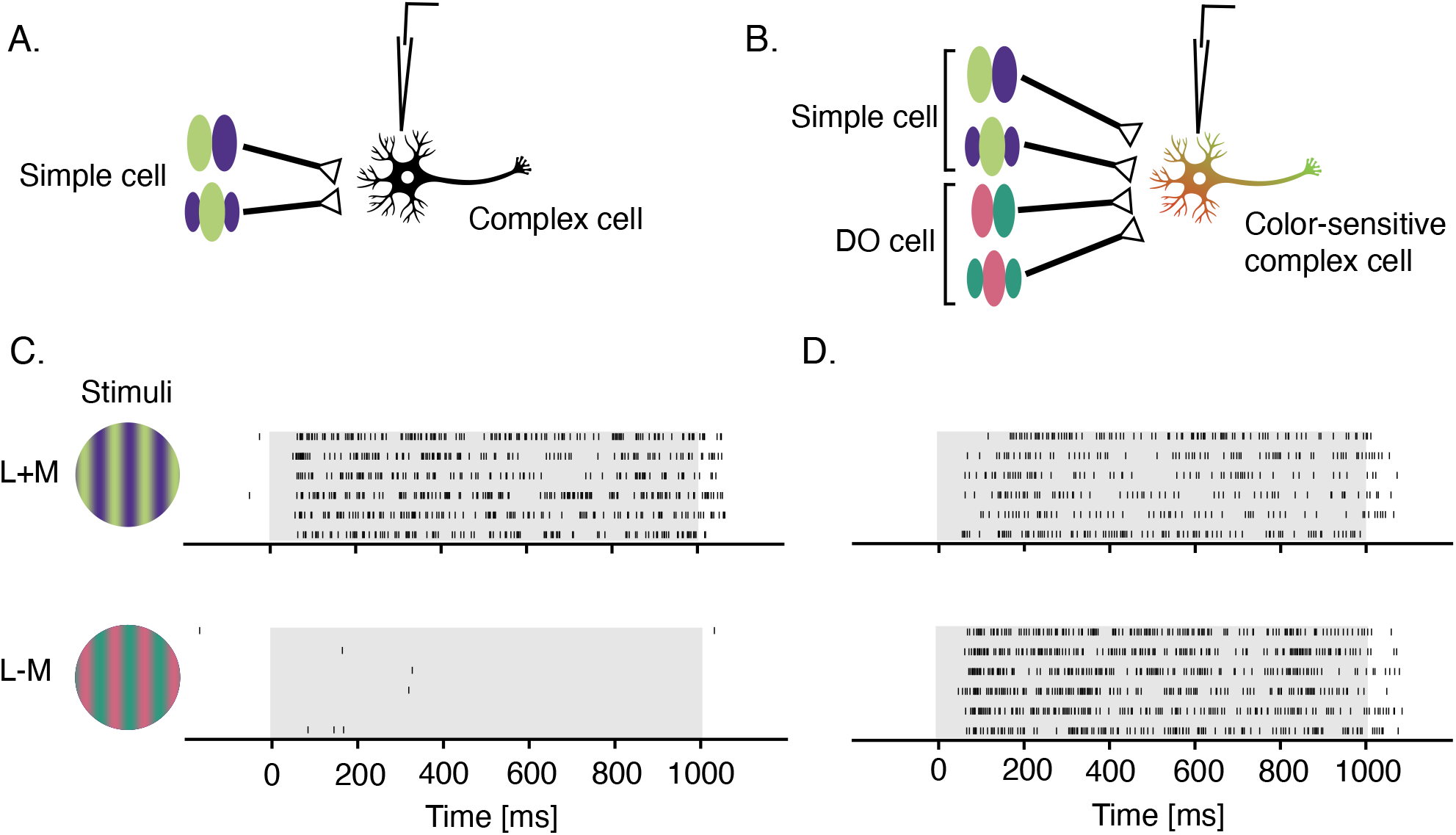
Proposed signal convergence of simple cells and DO cells onto complex cells **A.** A hypothetical complex cell receiving input from simple cells with overlapping odd- and even-symmetric receptive fields. **B.** A hypothetical color-sensitive complex cell receiving input from simple and DO cells. **C.** Response of a complex cell to a drifting sinusoidal grating that modulates L- and M-cones at 3 Hz with identical contrast in phase (top) and in anti-phase (bottom). **D.** Same as **C** but for a color-sensitive complex cell. Gray overlays indicate stimulus duration.

### Analogous neural coding of color and stereopsis

The stereotyped microcircuitry of area V1 contributes to vision for form, color, depth, and motion. These distinct visual modalities have distinct computational demands but V1 circuitry may contribute to each via a small set of operations that process different signals in similar ways. For example, parallels between the V1 processing of binocular disparity and motion direction are well established (Adelson and Bergen, 1991). We speculate that color and stereopsis have heretofore unappreciated parallels, and that models of binocular disparity tuning may provide a useful guide for the study of cone-opponent and non-opponent signal combination in V1.

A fundamental component of binocular V1 models are simple cells that linearly sum signals from the two eyes (Anzai et al., 1999b; Read and Cumming, 2003). Analogous building blocks in the color domain are simple cells and DO cells, which sum cone non-opponent and cone-opponent signals with a similar degree of linearity. Binocular simple cells are thought to provide input to binocular complex cells that implement an energy calculation (Anzai et al., 1999a; Read and Cumming, 2003). The analogous convergence of simple cell and DO cell outputs would implement a spectral energy calculation (Horwitz and Hass, 2012; Barnett et al., 2020).

The binocular energy model, while extremely successful in describing complex cell responses, requires refinement (Haefner and Cumming, 2008). For one thing, it fails to account for the attenuation of responses to anti-correlated signals between the two eyes (Cumming and Parker, 1997). This specialization of real V1 cells is thought to reflect the statistics of natural inputs to the visual system (Haefner and Cumming, 2008). Under natural viewing conditions, binocularly correlated patterns are more common than anticorrelated patterns, and V1 neurons appear specialized to encode them.

A parallel phenomenon may exist in the domain of color. Under natural viewing conditions, luminance and chromatic spatial gradients tend to be aligned, and their alignment (or misalignment) carries important information regarding the physical sources of the gradients. Edges between different materials under fixed illumination produce in-phase luminance and chromatic modulations, whereas uncorrelated variations in illumination and pigmentation, such as produced by curved 3-dimensional objects of non-uniform reflectance, produce out-of-phase modulations (Kingdom, 2003; Kunsberg et al., 2018). The energy model in **Figure 4B** produces phase-invariant responses and so does not account for these specializations. Whether real V1 neurons respond in accordance with the energy model or show enhanced responses to natural alignments between chromatic and luminance signals is an important and unanswered question.

In summary, DO cells and simple cells have much in common. As shown by the current study, both cell types combine signals roughly linearly across their RFs and, as shown previously, they share a Gabor-like RF structure (De and Horwitz, 2020). These observations motivate the idea that simple cells and DO cells are closely related neuronal types that may contribute similarly to downstream circuits that integrate cone-opponent and cone-non-opponent signals for spatial image analysis.

## MATERIALS & METHODS

### Contact for Resource Sharing

Further information and requests for resources should be directed to and will be fulfilled by the Lead Contact, Gregory D. Horwitz.

### Resources Table

| REAGENT or RESOURCE | SOURCE | IDENTIFIER |
| --- | --- | --- |
| **Experimental Models:**<br>**Organisms/Strains** |  |  |
| Rhesus monkeys (*Macaca Mulatta*) | Washington National Primate Research Center | N/A |
| **Software and Algorithms** |  |  |
| MATLAB | Mathworks | https://www.mathworks.com/products/matlab.html RRID: SCR_001622 |
| Plexon Sort Client | Plexon | http://www.plexon.com RRID: SCR_003170 |
| Plexon Offline Sorter | Plexon | http://www.plexon.com RRID: SCR_000012 |

### General

All protocols conformed to the guidelines provided by the US National Institutes of Health and the University of Washington Animal Care and Use Committee. Data were collected from two adult male rhesus macaques (Macaca mulatta). Each monkey was surgically implanted with a titanium headpost and a recording chamber (Crist Instruments) over area V1. Eye position was monitored continuously using either an implanted monocular scleral search coil or a digital eye-tracking system (SMI iView X Hi-Speed Primate, SensoMotoric Instruments).

### Task

The monkeys sat in a primate chair 1 m from a cathode ray tube (CRT) monitor (Dell Trinitron Ultrascan P991) in a dark room during the experiments. In a subset of sessions, the distance was reduced to 0.7 m and the pixel size was changed accordingly to preserve angular subtense. During white noise presentation, the monkeys fixated a centrally located dot measuring 0.2 x 0.2° and maintained their gaze within a 1.6 x 1.6° fixation window. During the closed-loop isoresponse measurements, the monkeys maintained their gaze within a 0.8 x 0.8° window. Successful fixation was rewarded with apple juice, and fixation breaks aborted trials.

### Monitor Calibration

Monitor calibration routines were adapted from those included in Matlab Psychophysics toolbox (Brainard, 1997; Pelli, 1997). The emission spectrum and voltage-intensity relationship of each monitor phosphor were measured with a spectroradiometer (PR650, PhotoResearch Inc.). Stimuli were gamma-corrected in software to compensate for the non-linearity of these voltage-intensity relationships. The color resolution of each channel was increased from 8 to 14 bits using a Bits++ video signal processor (Cambridge Research Systems, Ltd.). The monitor refreshed at 75 Hz and background was uniform gray (x = 0.3, y = 0.3, Y = 55–75 cd/m^2^).

### Electrophysiological recordings

We recorded from well-isolated V1 neurons (RF eccentricity: 1.3°–5.9°, median = 3.6°) using extracellular tungsten microelectrodes (Frederick Haer, Inc.) lowered through the dura mater via hydraulic microdrive (Stoelting Co.). Electrical signals were amplified, digitized at 40 kHz (Plexon, Inc.), and recorded.

### Experimental Protocol

Each experiment consisted of three phases. During the first phase, spatiochromatic tuning was probed with a white noise checkerboard stimulus and data were analyzed online by spike-triggered averaging. During the second phase, the white noise stimulus was customized to the RF of each neuron. During the third phase, high-contrast images with the same spatial structure used in *Phase 2* were presented for relatively long durations (300 ms). Each of these phases is detailed below.

### Phase 1: Checkerboard white noise

Each stimulus frame contained a 10 x 10 grid of pixels each of which subtended 0.2 x 0.2° (**Figure 2A**) (Horwitz et al., 2007). The stimulus changed every 13.33 ms. The intensity of each phosphor at each pixel was modulated independently according to a Gaussian distribution with a standard deviation of 15% of the physically achievable range. The space-time averaged intensity of each phosphor was equal to its contribution to the background.

Neuronal responses to white noise stimuli were analyzed by spike triggered averaging (**Figure 2A**). In this analysis, the 15 frames preceding every spike were collected and averaged across spikes. From these 15 spike-triggered average (STA) frames, we selected online the one that differed most from the background and identified pixels that differed significantly from the background (p<0.05, z-tests performed on each phosphor separately). These data were used to customize the white noise stimulus to the RF in *Phase 2* of the experimental protocol (see below). We selected for additional study only cells whose RFs consisted of at least two subfields with distinct chromatic preferences (**Figure 2A**).

The checkerboard white noise stimulus modulated neurons weakly for three reasons. First, individual stimulus pixels were small relative to V1 RFs. This was necessary to distinguish one RF subfield from another but resulted in each subfield being stimulated by independent pixel modulations that tended to cancel. Second, the pixels modulated rapidly, so multiple frames were effectively averaged together in the early visual system, prior to V1. Third, phosphor intensities were drawn from Gaussian distributions. Most of the probability mass of a Gaussian distribution is near the mean, which was identical to the background, so high contrast pixels were improbable (**Figure S1A–B**).

### Phase 2: Hyperpixel white noise

For each neuron with an STA containing at least two spatially distinct subregions (with distinct chromatic preferences), we created a custom white noise stimulus by yoking the pixels within each of the two subfields (**Figure 2B**). Phosphor intensities at the two yoked collections of pixels (the two hyperpixels) were modulated according to the same Gaussian distribution used in *Phase 1.* Pixels outside of the RF were not modulated.

To examine how signals were combined across the two targeted subfields, we represented hyperpixel stimulus frames as a six-dimensional vectors of background-subtracted RGB values and then projected each of these segment of vectors onto the two halves of the temporo-chromatic STA. This operations produces two scalar values that indicated how strongly short, overlapping segments of the stimulus movie drove the two RF subfields. We visualized a firing rate map from these projections by computing the ratio of spike-triggered stimuli to the total stimuli (Chichilnisky, 2001). We also fit the data with linear and non-linear models and compared the fits to examine spatial integration quantitatively. The details of this analysis can be found in the section: **White noise analysis of signal combination across subfields**.

### Phase 3: Isoresponse measurement

We selected the STA frame that differed most from the background and separated it into its two hyperpixels, each of which selectively stimulated one RF subfield with its preferred light (represented along the 45° and 135° directions in **Figure 3A**). We then linearly combined these two images in different proportions to create a family of stimuli that can be represented in the same plane used to construct the firing rate map in *Phase 2* **(Figure 2C)**. The origin of the coordinate system represents the gray background of the display. Direction represents the overall contrast between the two halves of the stimulus, and distance from the origin represents stimulus contrast relative to the background. No universally accepted definition of contrast applies to all color directions. Therefore, for convenience, we defined contrast along the 45° direction as the projection of the RGB values onto one half of the STA. Contrast along the 135° direction was defined similarly using the other half of the STA.

### Contrast staircase procedure

To examine interactions between subfields, we identified collections of stimuli described above that evoked the same number of spikes using the following procedure. On each trial, the computer presented a stimulus and counted spikes from the onset response latency, defined as the peak frame of the STA from *Phase 2,* until the end of the stimulus presentation. This spike count was compared to an experimenter-defined target response (**Figure S3A**). If the spike count was lower than the target response, the contrast of the image was increased by a factor of 1.35. If the spike count exceeded the target response, the contrast of the image was decreased by a factor of 0.65. This process continued until a reversal occurred. A reversal is a response that exceeded the target response after having fallen below it on the previous stimulus presentation or a response that fell below the target having exceeded it on the previous stimulus presentation. After each reversal, the change in contrast per trial decreased by 75% (**Figure S3B**). The staircase halted after seven reversals or whenever the contrast exceeded the physical limitations of the display. Staircase termination points were taken as estimates of the contrast that evoked the target response. Presentations of stimuli in pairs of directions in the stimulus space were randomly interleaved to mitigate non-stationarity due to adaptation. Each stimulus was presented for 300 ms and was separated from the preceding and subsequent stimuli by more than 1 s.

### Cell Screening

We recorded from 232 well-isolated V1 neurons and made isoresponse measurements from 98 of them. Neurons were classified as “simple”, “double-opponent” or “NSNDO” (neither simple nor double-opponent) on the basis of responses to white noise as described below.

### Cone weights

Cone weights were calculated from *Phase 2* of the experimental protocol. For each cell, we identified the STA frame that differed maximally from the background and computed a weighted average of this frame and the two flanking frames. The weight of each frame was proportional to the sum of squared red, green and blue intensities relative to the background. We decomposed this weighted STA by singular value decomposition into a color weighting function and a spatial weighting function, defined as the first row and column singular vectors, respectively (De and Horwitz, 2020). Together, the color and spatial weighting functions captured 96.7±5.0% (mean±SD) of the variance in the weighted STAs. The color weighting function was converted to cone weights that are assumed to act on cone contrast signals (Weller and Horwitz, 2018). The spatial weighting function of every cell consisted of one positive and one negative weight, because only neurons with spatially opponent RFs were recorded.

### Spike-triggered covariance analysis

Cone weights are interpretable only under a linear model of signal combination across cone types (Weller and Horwitz, 2018). One way to test this assumption is by analysis of spike-triggered covariance. To implement such a test, we computed the first principal component (PC1) of the spike-triggering stimuli orthogonal to the STA (Touryan et al., 2002; Horwitz et al., 2005; Rust et al., 2005). A PC1 that is larger than expected by chance reveals a nonlinear component of the cell’s response to the white noise stimulus. We assessed the significance of the PC1 by randomly shifting spike trains in time relative to the *Phase 2* stimulus movie, recalculating the PC1, and obtaining its eigenvalue (Rust et al., 2005). This procedure was repeated 1000 times. If the largest eigenvalue from the unrandomized data exceeded 95% of the largest eigenvalues from the randomized data sets, we concluded that the PC1 was significant at the 0.05 level. Neurons with a significant PC1 were classified as NSNDO.

Neurons lacking a significant PC1 were classified as simple if their L- and M-cone weights had the same sign, accounted for 80% of the total cone weight and individually accounted for at least 10%. None of the simple cells we studied showed evidence of opponent input from the S-cones, but some appeared to receive a small non-opponent S-cone input. Twenty-six cells in our data set were categorized as simple. Our criteria for DO cells were a lack a significant PC1 and a pair of cone weights of opposite sign. To ensure that all DO cells were truly cone-opponent, weights of small absolute value were ignored; to be classified as DO, a neuron had to have an S-cone weight that accounted for at least 20% of the total or L- and M-cone weights that accounted for at least 80% jointly and 20% individually. Twenty-five cells were categorized as DO. The 47 neurons that did not meet the DO or simple cell criteria were classified as NSNDO. These criteria are arbitrary but the central results of this study are robust to these particulars (**Figure S5**). We describe the cell classification criteria below that was used for obtaining the results in **Figure S5**.

### Luminance tuning index

A luminance tuning index was obtained by projecting the normalized cone weights of each cell to the cone weights of a luminance mechanism. The luminance cone weights were estimated by regressing the Stockman-Macleod-Johnson 2° cone fundamentals onto the Judd-Vos 1978 2° photopic luminosity function to find the best-fitting coefficients (0.83 L + 0.55 M + 0.03 S) (Vos, 1978; Stockman et al., 1993). The luminance tuning index ranged from 0 to 1. Cells were classified as DO if their index value was < 0.33 and they lacked a significant PC1. Twenty-two cells were classified as DO. Cells were classified as simple if their index value was > 0.67 and they lacked a significant PC1. Twenty-seven cells were classified as simple. The remaining 49 neurons were classified as NSNDO.

### White noise analysis of signal combination across subfields

We fit the data from *Phase 2* of the experimental protocol with a generalized linear model (GLM) and a generalized quadratic model (GQM).

The GLM was defined as:

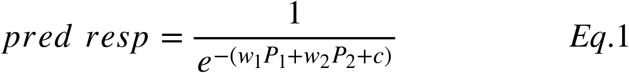

where *pred resp* is the predicted response of the neuron and *P*_1_ and *P*_2_ are the projection magnitudes of the short segments of the stimulus movie onto the two halves of the hyperpixel STA. *w*_1_, *w*_2_, and *C* were fit using the MATLAB routine (*fitglm*) to maximize the binomial likelihood of a spike.

The GQM was defined similarly as:

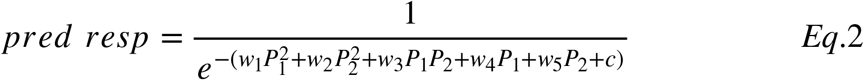

### Evaluating the performance of generalized linear and quadratic models

We quantified the ability of the fitted models to predict whether or not each stimulus segment evoked a spike using receiver operating characteristic (ROC) analysis (Green and Swets, 1966). Classification error was defined as 1 minus the area under the ROC curve (**Figure S2B**). To avoid overfitting, the model was fit with 90% of the data and tested on the remaining 10%. The white noise non-linearity index (white noise NLI) for each cell was defined as:

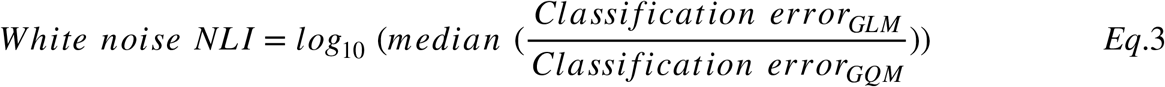

where the median is taken is taken across 10 cross-validation data partitions.

### Model fits to isoresponse staircase termination points

To assess the linearity of signal integration across the gamut of our video display, we fit the staircase termination points from *Phase 3* with linear and quadratic models. Fitting was performed using a standard inbuilt MATLAB routine for function minimization (*fmincon*) to minimize the Tukey-bisquare objective function (Fox, 2002).

Searches for the stimuli that produced the target response were conducted in multiple directions of the stimulus space (e.g. **Figure 3A–C**), but angles were fixed. We therefore fit the data with a model that assumes radial error. The linear model can be written as:

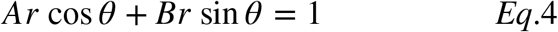

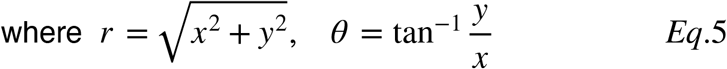

*x* represents the projection of each image onto one hyperpixel of the STA and *y* represents the projection onto the other hyperpixel. *A* and *B* are fitted parameters.

The quadratic model can be written as:

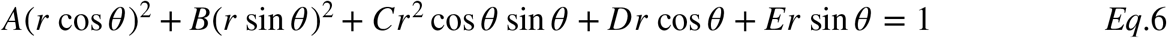

Where *A*, *B*, *C, D* and *E* are fitted parameters.

### Evaluating model fits to staircase termination points

We evaluated the quality of model fits by calculating the sum of Tukey-bisquared errors between the data and the model predictions. To avoid overfitting, we used leave-one-out cross validation. The isoresponse non-linearity index (isoresponse NLI) was defined as the median of the ratio of cross-validated linear model errors and quadratic model errors in logarithmic units.

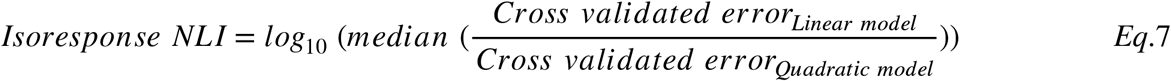

### Drifting gratings

Two neurons were stimulated with drifting, sinusoidal gratings (2 cycles per degree, 3 Hz, 1° diameter circular aperture) that modulated the L- and M-cones with identical contrasts either in phase (L+M) or in anti-phase (L-M) (**Figure 4**). These neurons were not tested with the standard protocol.

## SUPPLEMENTARY FIGURE LEGENDS

**Figure S1.**
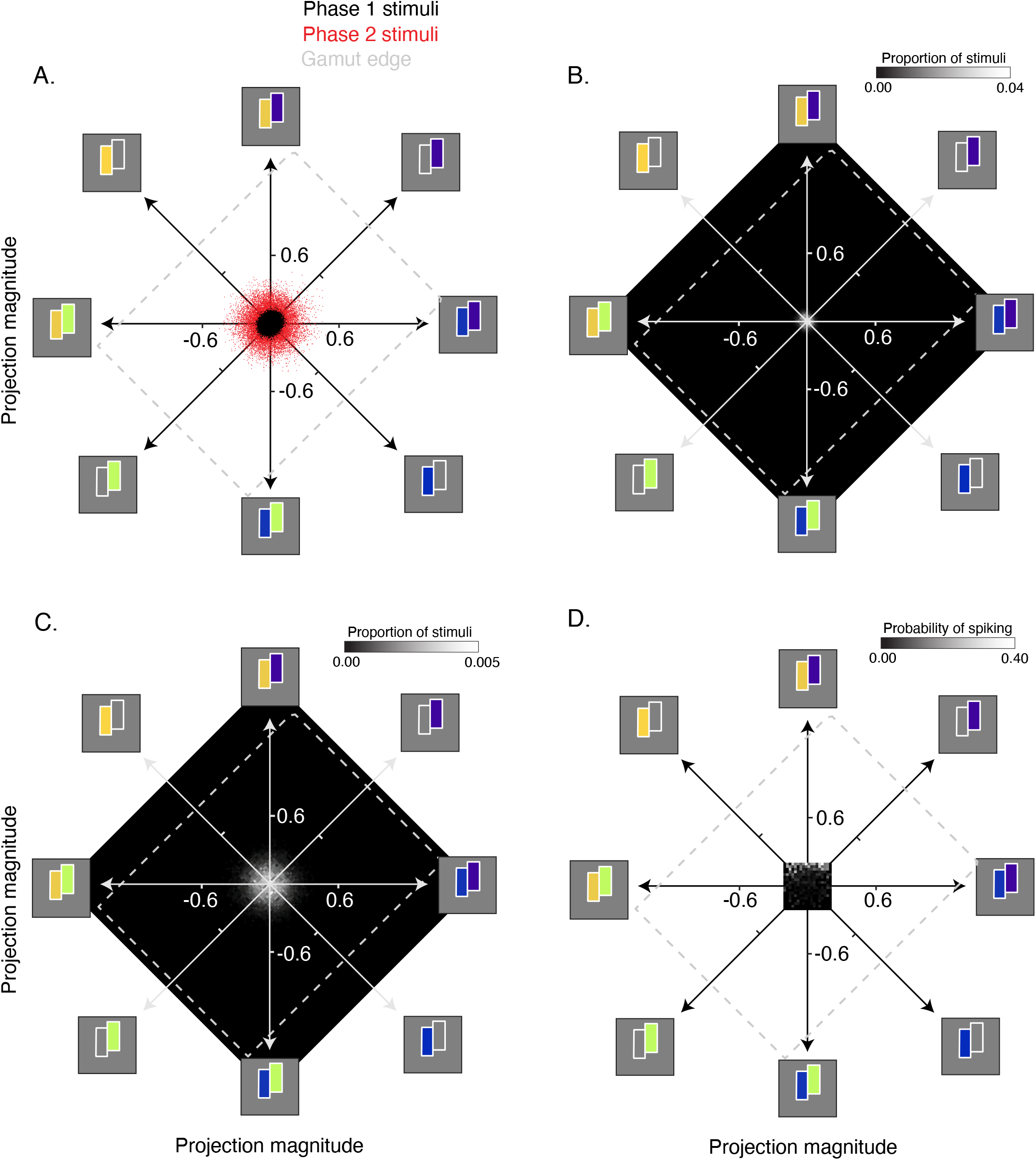
Comparison of the effective stimulus contrast in the three phases of the experiment. **A.** Stimuli from *Phase 1* (black) and *Phase 2* (red) were projected onto the spatial–temporal–chromatic STA shown in **Figure 2B.** Projection magnitudes of both stimuli occupy only a small region of the display gamut (dashed gray box). **B.** Twodimensional histogram of the *Phase 1* projections shown in **A**. **C.** Same as **B** but for the *Phase 2* stimuli. **D.** The probability of spiking as a function of hyperpixel stimulus projections onto the two halves of the STA. Projection magnitudes from the 5th to the 95th percentile are shown. Within this range, the probability of spike increases approximately as a a linear combination of the stimulus projections, but this range is a small fraction of what can be achieved on the display.

**Figure S2.**
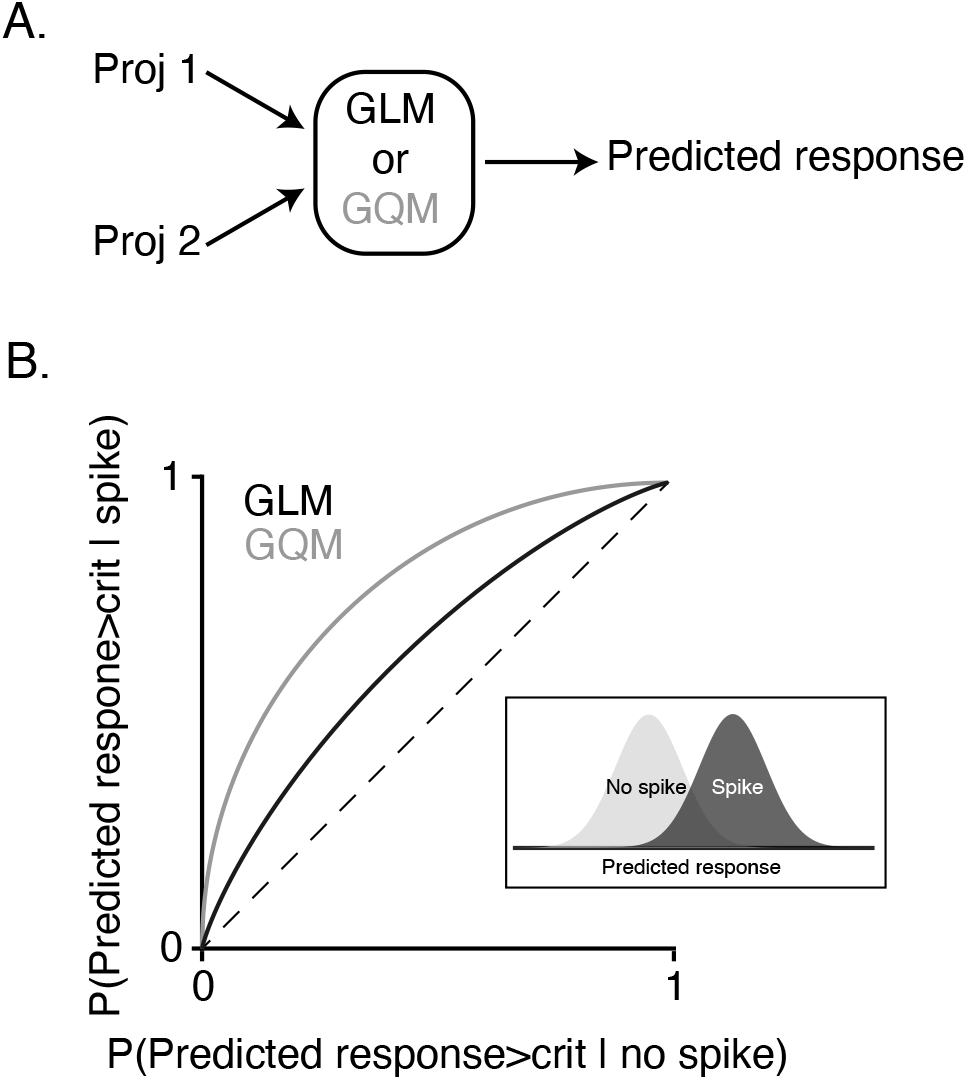
Analysis of neuronal spatial integration of white noise stimuli **A.** Probability of spiking was predicted as a function of projection magnitudes onto the two halves of the STA (Proj 1 and Proj 2) using a generalized linear model (GLM) or a generalized quadratic model (GQM). **B.** An ROC analysis was used to assess the ability of the GLM and GQM to classify stimuli into those that did not evoke a spike (inset, gray) and those that did (inset, black).

**Figure S3.**
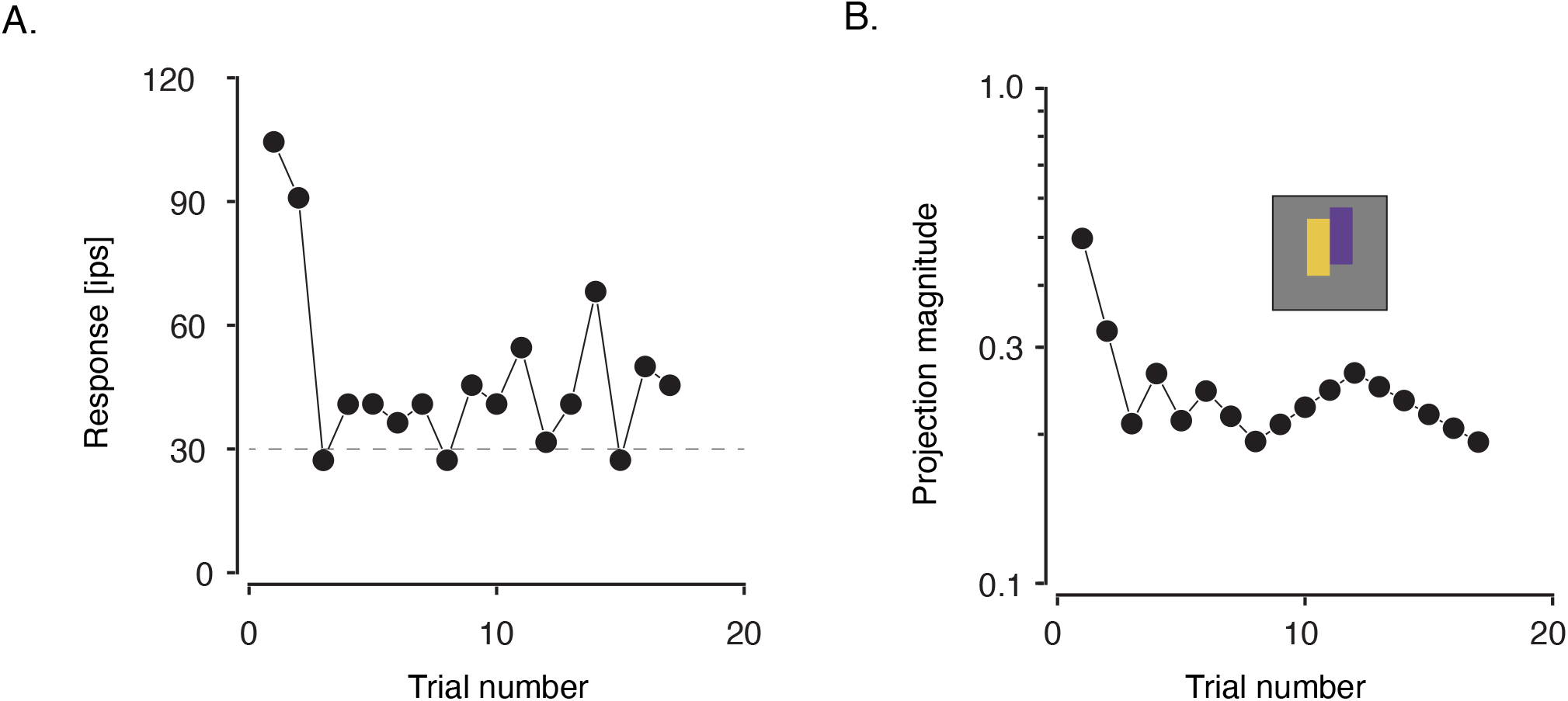
An example staircase from the closed-loop procedure used to study the DO cell in **Figure 2A.** Neuronal response (in impulses per second, ips) is plotted as a function of trial number (intervening stimuli skipped). The target firing rate was 30 ips (dashed line). **B.** The projection magnitude as a function of trial number for the same staircase. The staircase termination point is defined as the projection magnitude of the stimulus presented in the final (17^th^) trial.

**Figure S4.**
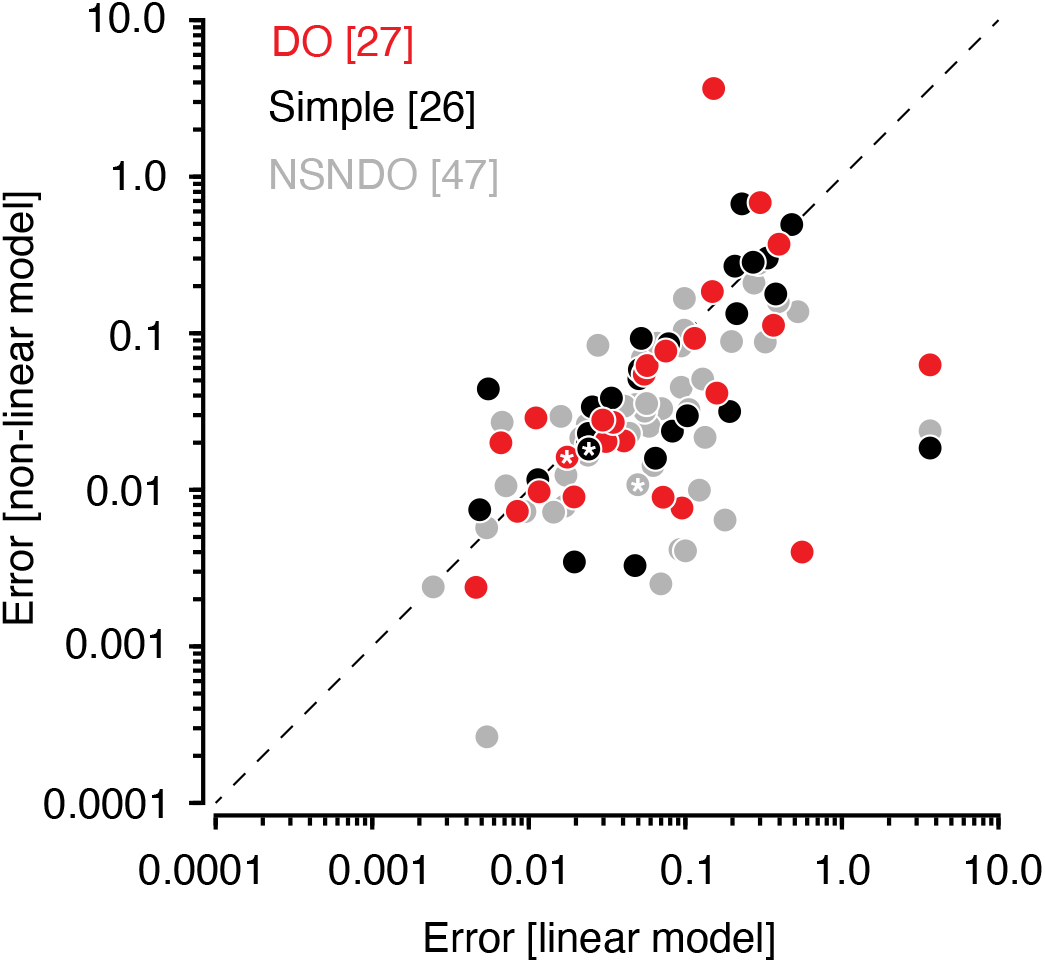
Cross-validated errors from linear and non-linear fits to data from *Phase 3* for DO cells (red), simple cells (black), and NSNDO cells (gray). Example neurons from **Figure 3** are marked with white asterisks.

**Figure S5.**
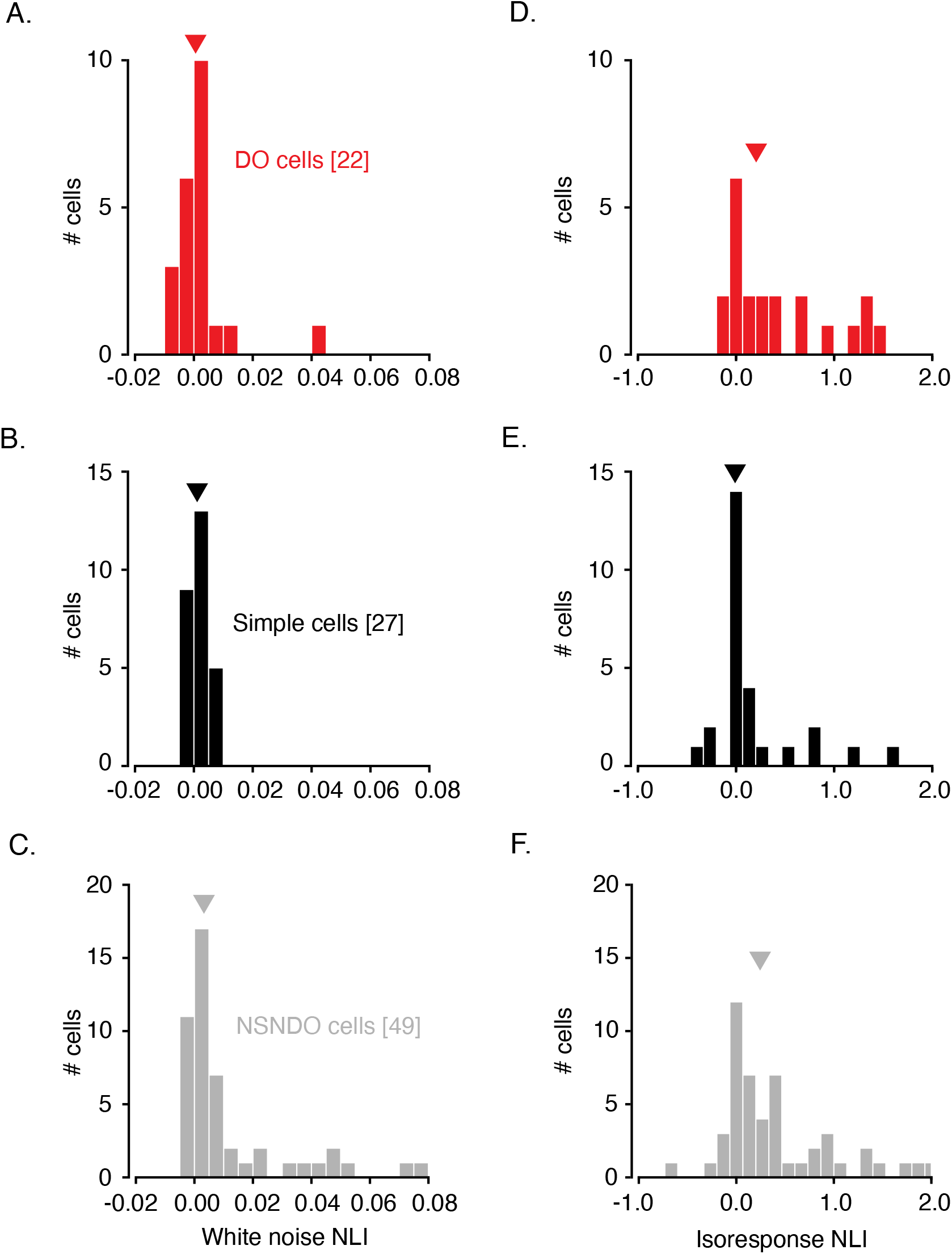
Reclassification of neurons with a different cone weight criteria. A luminance tuning index was calculated for each cell by weighting and summing normalized cone weights (0.83 L + 0.55 M + 0.03 S). This index ranges from 0 to 1. Cells were classified as DO if their index value was < 0.33 and they had an insignificant PC1. Cells were classified as simple if their index value was > 0.67 and they had an insignificant PC1. Remaining cells were classified as NSNDO. Histograms of white noise NLIs for DO (**A**), simple (**B**) and NSNDO (**C**) cells classified this way. White noise NLIs of DO and simple cells were similar (p=0.78, Mann-Whitney U Test), and were lower than NSNDO neurons (p=0.004, Mann-Whitney U test). **D–F.** Identical to **A–C.** but showing isoresponse NLIs. Isoresponse NLIs of DO and simple cells were lower than NSNDO neurons (p=0.06, Mann-Whitney U test).

## ACKNOWLEDGEMENTS

We thank Yasmine El-Shamayleh, Fred Rieke, Greg Field and Jacob Yates for comments on the manuscript. This work was funded by NIH EY018849 to Gregory D. Horwitz, NIH/ORIP grant P51OD010425, and NEI Center Core Grant for Vision Research P30 EY01730 to the University of Washington and R90 DA033461 (Training Program in Neural Computation and Engineering) to Abhishek De.

